# Microbial DNA on the move: sequencing based detection and analysis of transduced DNA in pure cultures and microbial communities

**DOI:** 10.1101/2020.01.15.908442

**Authors:** Manuel Kleiner, Brian Bushnell, Kenneth E. Sanderson, Lora V. Hooper, Breck A. Duerkop

## Abstract

Horizontal gene transfer (HGT) plays a central role in microbial evolution. Our understanding of the mechanisms, frequency and taxonomic range of HGT in polymicrobial environments is limited, as we currently rely on historical HGT events inferred from genome sequencing and studies involving cultured microorganisms. We lack approaches to observe ongoing HGT in microbial communities. To address this knowledge gap, we developed a DNA sequencing based “transductomics” approach that detects and characterizes microbial DNA transferred via transduction. We validated our approach using model systems representing a range of transduction modes and show that we can detect numerous classes of transducing DNA. Additionally, we show that we can use this methodology to obtain insights into DNA transduction among all major taxonomic groups of the intestinal microbiome. This work extends the genomic toolkit for the broader study of mobile DNA within microbial communities and could be used to understand how phenotypes spread within microbiomes.

**Significance Statement:** Microbes can rapidly evolve new capabilities by acquiring genes from other organisms through a process called horizontal gene transfer (HGT). HGT occurs via different routes, one of which is by the transfer of DNA carried by microbe infecting viruses (phages) or virus-like agents. This process is called transduction and has primarily been studied in the lab using pure cultures or indirectly in environmental communities by analyzing signatures in microbial genomes revealing past transduction events. The transductomics approach that we present here, allows for the detection and characterization of genes that are potentially transferred between microbes in complex microbial communities at the time of measurement and thus provides insights into real-time ongoing horizontal gene transfer.

## Introduction

The importance of horizontal gene transfer (HGT) as a driver of rapid evolution and adaptation in microbial communities and host-associated microbiomes has become increasingly recognized(1, 2). Publicly available genomes and metagenomes have revealed pervasive horizontally acquired genes in almost all available genomes. A study of HGT in the human microbiome, for example, showed >10,000 recently transferred genes in 2,235 analyzed genomes(3). HGT has been implicated in the spread of antibiotic resistance genes(4), toxin and other virulence genes(5, 6), as well as genes that enable digestion of dietary compounds by microbes in the intestine(7), and metabolic genes that augment microbial metabolism with critical functions in environmental populations(8). Despite its recognized importance, our understanding of the taxonomic range, frequency, and mechanisms of HGT are still limited. Most studies of HGT in microbiomes rely on analysis of microbial genomes(3, 9) and as such these methods attempt to reconstruct historical HGT. What we currently lack are methods that measure ongoing HGT and identify the mechanism of DNA transfer. Here we present a novel method that specifically determines the sequence of DNA that is transferred between cells via one of the major known pathways for DNA transfer – transduction.

Currently, there are three major ways that genetic material is known to be exchanged between microbial cells, (1) transformation – uptake of DNA by naturally competent cells, (2) conjugation – exchange of genetic material (e.g. plasmids) using direct contact between donor and recipient cells, and (3) transduction – transfer of genetic material by viruses or virus-like particles (VLPs)(2). Here we focus on transduction only. There are several known types of transduction including classic specialized and generalized transduction, and more recently discovered types, including gene transfer agents (GTAs), lateral transduction and hijacking of bacteriophage (phage) particles by genomic islands(10–12). During specialized transduction DNA adjacent to prophage integration sites in the bacterial genome are co-excised at a low frequency and packaged into phage heads after prophage genome replication. In generalized transduction non-random pieces of the host bacterial genomic DNA or plasmids get packaged at low frequency into phage particles when a lytic phage infects and replicates in a bacterial cell. This non-random packaging is mediated by genomic features that resemble the packaging site (*pac* site) on the phage genome, which is used by the phage particle packaging machinery as the start site phage DNA packaging into the capsid(13). In lateral transduction prophages replicate while still integrated in the host genome and prophage packaging initiates *in situ* ultimately leading to high frequency packaging of host DNA in a unidirectional fashion away from the prophage integration site(12).GTAs are phage-like particles encoded in bacterial genomes that package random pieces of the genomic DNA upon production and can transfer these pieces to other cells(10). In contrast to phages, GTAs do not carry the DNA content sufficient to support their reproduction in the target cells. Lastly, some genomic islands, including pathogenicity islands, can hijack phages capsids in an act of molecular piracy that enables their transduction(14, 15).

The unifying characteristic of all types of transduction is that virus or VLPs serve as the vector for transfer of genetic material between cells. Evidence so far indicates that these particles are abundant in most environments and that transduction occurs with a high frequency(16, 17). However, approaches for measuring the abundance of transducing particles and transduction frequencies in microbiome samples are limited. These approaches usually rely on the application of cultured phage to environmental samples(16, 17) or sequencing of bacterial 16S rRNA genes from purified VLPs(18). The latter approach can determine which bacterial taxa’s DNA is carried in a VLP. However, the approach is limited to marker genes for which conserved PCR primer pairs exist and thus the majority of transduced DNA cannot be detected.

Here we describe an unbiased approach, termed “transductomics”, which uses DNA sequencing to identify and characterize DNA originating from microbial cells that is carried in VLPs. This DNA is thus part of the pool of potentially transduced DNA termed the “transductome”. Our approach is based on two observations of transduced DNA in VLPs. First, transduced DNA often represents the genome of hosts that are present in the same sample as the VLPs. Therefore, if DNA from a microbe is found within VLPs purified from the same sample this indicates a potential transduction event. Second, unique regions of the microbial host’s genome are unevenly enriched in the VLPs, as most mechanisms of transduction do not lead to random packaging of the host’s genome. In recent years, the uneven sequence coverage patterns produced by phages or GTAs carrying microbial host DNA have been used to characterize the genome biology and mechanisms of DNA packaging of specific host-phage/GTA systems(19–22). Our transductomics approach exploits these sequence coverage patterns to identify and characterize transduced DNA in microbial communities. In the past, host DNA carried by VLPs may have been sequenced during metagenomic sequencing of purified VLPs, but without appropriate analysis tools these host derived sequences were classified as host contamination of the VLP sample rather than being recognized as transduced DNA(23).

The transductomics approach that we present requires the sequencing of both the complete microbial community sample, and VLPs that are ultra-purified using CsCl density gradient centrifugation from the same sample (Fig. 1). The VLP and complete sample sequencing reads are mapped to long genome contigs assembled from the complete sample metagenome. These contigs represent both microbial and viral genomes. Visualization of the read mapping coverages along the contigs comparing VLP and complete metagenome read coverages reveals patterns that can be associated with host DNA transport via VLPs. We demonstrate this method first using pure culture models of different transducing phages and other transducing particles. This is followed by the application of the approach to a murine intestinal microbiome community.

**Figure 1:**
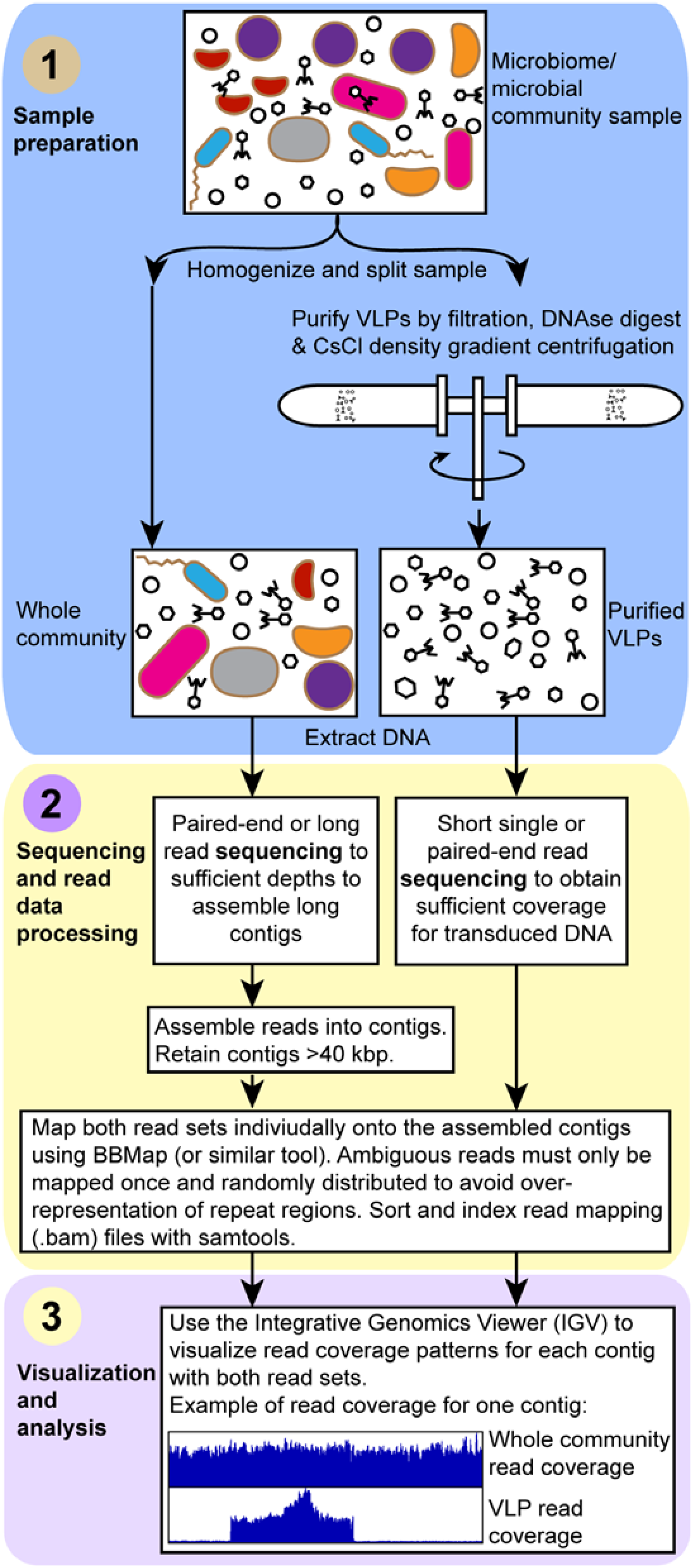
The “transductomics” workflow. In the sample preparation step the sample is gently homogenized and split into two subsamples. One subsample is directly used for whole community DNA extraction, the other subsample is subjected to ultra-purification of virus-like particles (VLPs) using a combination of filtration, DNAse digest and CsCl density gradient centrifugation as previously described(24) followed by DNA extraction from the purified VLPs. Both DNA samples are sequenced to different depths and potentially with different sequencing approaches, although in many cases the same sequencing approach could be applied to both samples. For the whole community DNA sample, the sequencing must focus on ultimately achieving assembly of long metagenomics contigs. For the VLP DNA sample, the sequencing must focus on maximal read coverage, and no assembly is needed for these reads. The whole community sequencing reads are assembled using a suitable assembler. Contigs smaller than 40 kbp are discarded. Both the whole community and VLP sequencing reads are mapped onto the contigs >40 kbp using BBMap(25) ensuring that ambiguously mapped reads are only used once and randomly assigned. To find transduced regions, the contigs read coverage patterns for both whole community and VLP reads are visualized using the Integrative Genomics Viewer(26).

## Results and Discussion

### Characterization of sequence coverage patterns associated with different transduction modes in model systems

#### Specialized transduction by *Escherichia coli* prophage λ(27)

We used the well-studied specialized transducing bacteriophage λ to analyze the sequencing coverage patterns produced by specialized transduction. In specialized transduction a prophage, which is integrated in the chromosome of the bacterial host, packages host genome derived DNA with low frequency due to imprecise excision from the genome upon prophage induction. Prophage λ integrates between the *gal* (galactose metabolism) and *bio* (biotin metabolism) operons in the *E. coli* genome. In rare cases λ excision is imprecise and either the *gal* or the *bio* operon is excised and packaged in the phage particle (Fig. 2a)(27). This packaging of adjacent *E. coli* host derived DNA can lead to the transduction of the *bio* and the *gal* operons. Transduction of recipient cells can be temporary or permanent, depending on if the DNA gets recombined into the chromosome or remains as an extrachromosomal element, which is diluted out in the population during cell divisions.

**Figure 2:**
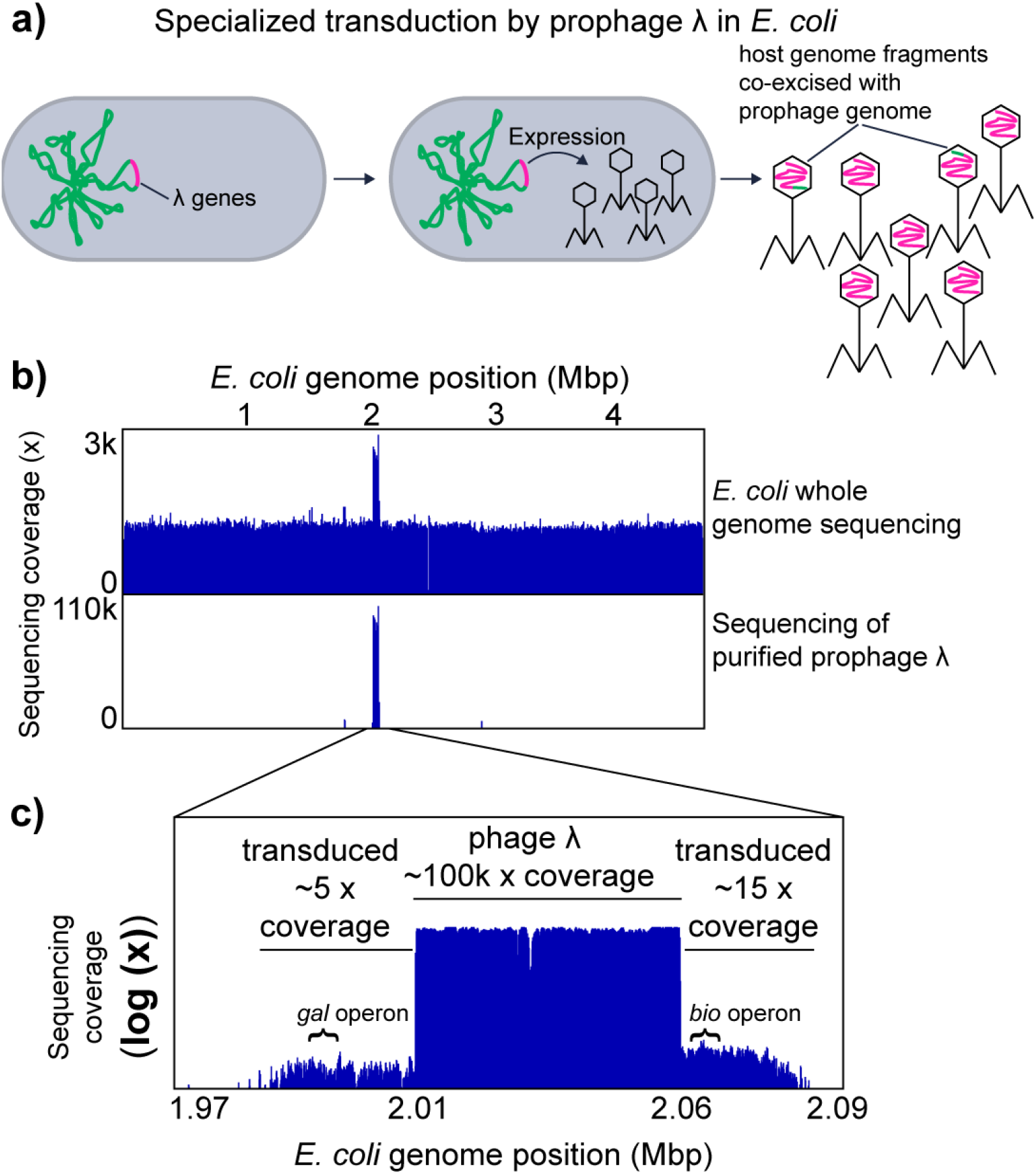
Specialized transduction by *E. coli* prophage λ. a) Illustration of specialized transduction. The prophage λ genome is integrated into the host chromosome. Upon induction of the prophage, the prophage genome is excised and replicated. The phage structural genes are expressed, phage particles are produced and the replicated phage genome is packaged into phage heads. Ultimately the phages are released to the environment by lysis of the host cell. Imprecise excision of the prophage λ genome happens at low frequency and leads packaging of the host chromosome into phage heads. These parts of the host chromosome can be transferred to new host genomes in the process called specialized transduction. b) Genome coverage pattern associated with prophage λ induction and specialized transducing prophage λ. The upper box shows coverage patterns for whole genome sequencing reads and purified phage particle reads mapped to the *E. coli* genome. c) In the lower box, an enlargement of the purified phage read coverage for the prophage λ region is shown (log scale). The positions of the *gal* and *bio* operons, which are known to be transduced by prophage λ, are indicated(27).

Using the transductomics approach we found that coverage of the *E. coli* genome with sequencing reads derived from purified λ phage particles is almost exclusively restricted to the λ phage integration site and two ~25 kbp regions on the left and right of the λ integration site (Fig. 2b and c). These flanking regions with read coverage represent the regions that are transduced by λ phage as indicated by the presence of the *bio* and the *gal* operons in these flanking regions (Fig. 2c). The coverage of the λ prophage region is roughly 10,000 fold greater than the coverage of the flanking transduced regions indicating that only a small number of phage particles actually carry transduced DNA and thus are specialized transducing particles.

Using the *E. coli*-prophage λ model we show that specialized transduction by a prophage produces a unique read coverage pattern. Furthermore, analysis of the read coverage pattern of the transduced DNA region adjacent to the prophage DNA allows determination of both the size and content of the transduced host genome region (~50 kbp in total in case of λ), as well as estimation of the frequency with which transducing particles are produced (1:10,000 in case of λ). The number of transducing particles produced based on our data is roughly 100-fold higher than previously reported values for successful transduction of the *gal* operon by phage λ (1:1,000,000 successful transductions per λ particles)(28), which indicates that only a small fraction of λ carrying host DNA ultimately leads to successful transduction.

#### Generalized transduction of the *Salmonella enterica* serovar typhimurium LT2 genome by phage P22 and the *E. coli* genome by phage P1

We used two well-studied generalized transducing bacteriophages P22 and P1 to analyze the sequencing coverage patterns produced during generalized transducing events. In generalized transduction nonspecific host chromosomal DNA is packaged into phage particles during lytic infection and can then be injected into a new host cell (Fig. 3a). The DNA can then recombine into the host chromosome by homologous recombination.

**Figure 3:**
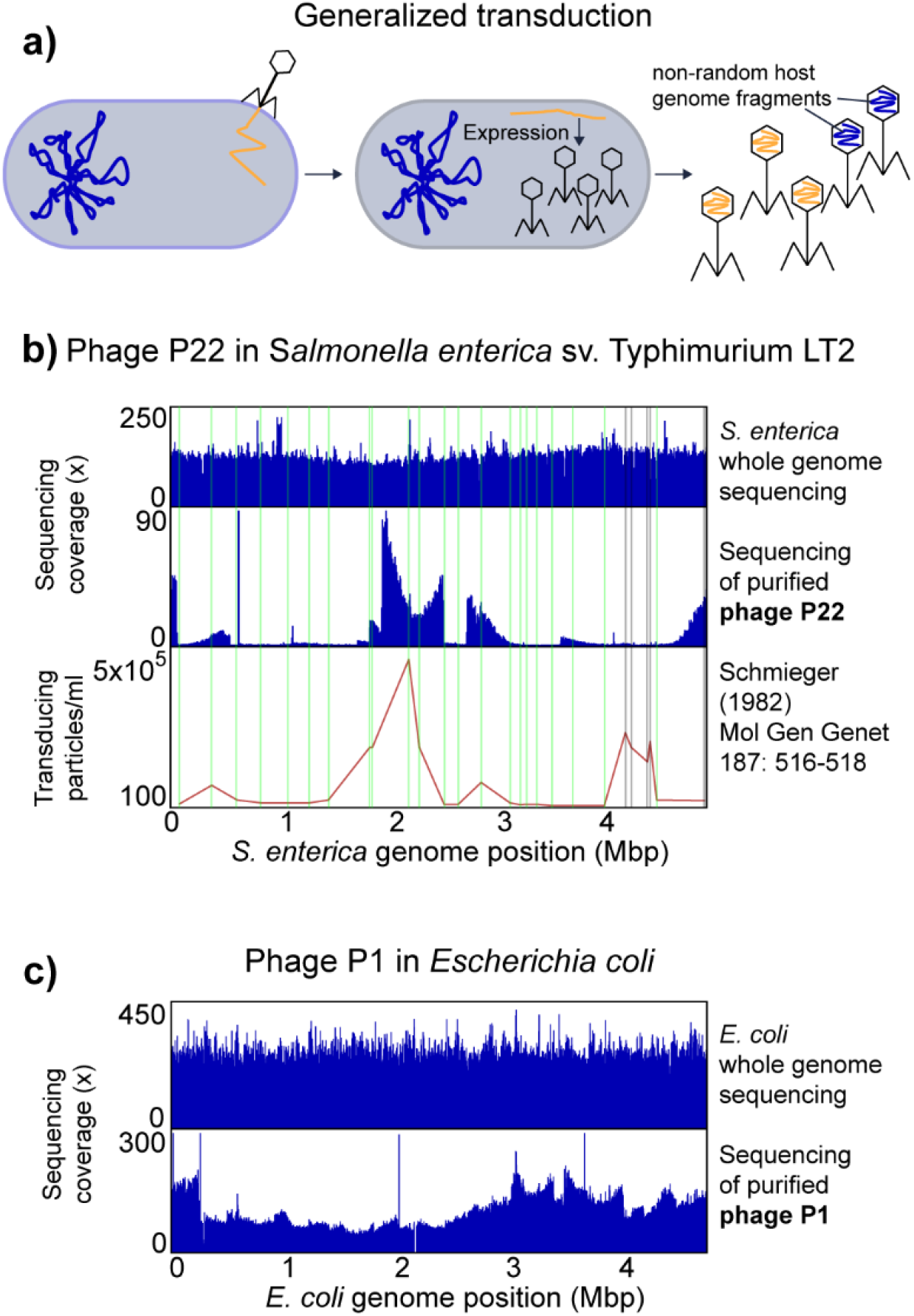
Generalized transduction by *S. enterica* phage P22 and *E. coli* phage P1. a) Illustration of generalized transduction. Upon phage infection, the phage genome is replicated in the host cell by rolling circle replication resulting in genome concatamers and phage particles are produced. The phage genome is packaged into the phage head by a so called head-full packaging mechanism, which relies on the recognition of a packaging (*pac*) site. The bacterial host chromosomes contain sites that resemble the *pac* site and thus lead to packaging of non-random pieces of the host chromosome into phage heads. The packaging happens in a processive fashion i.e. after one phage head has been filled the packaging machinery continues to fill the next phage head with the remaining DNA molecule. The likelihood that the packaging machinery dissociates from the molecule increases the further away from the *pac* site it gets, thus leading to a decreased packaging efficiency over distance. b) *Salmonella enterica* genome coverage pattern associated with generalized transduction by phage P22. Whole genome sequencing reads and purified phage particle reads were mapped to the *S. enterica* genome. In the lower part transduction frequencies for 28 chromosomal markers along the chromosome are shown as determined by Schmieger (1982)(30). Vertical lines indicate the positions of the chromosomal markers in green where the transduction frequency matches the read coverage, in grey where read coverage does not correspond to reported transduction frequency. c) *Escherichia coli* genome coverage pattern associated with generalized transduction by *E. coli* phage P1.

98.2% of the sequencing reads from purified P22 particles mapped to the P22 genome leaving 1.8% of reads that map to the *S. enterica* genome. The percentage of P22 particles mapping to the *S. enterica* genome corresponds to the reported percentage of 1.5% transducing P22 particles (i.e. carry host DNA) previously reported(29). The mapped P22 derived reads covered the *S. enterica* genome unevenly, while whole genome sequencing of *S. enterica* yielded even coverage (Fig. 3b). Regions of high or low P22 read coverage corresponded in 23 out of 28 previously reported transduced chromosomal markers(30) (Fig. 3b). Only one region at around 4 Mbp, for which high transduction frequencies had been reported, did not show high coverage (Fig. 3b), which might be due to differences in *pac* sites within this region between the *S. enterica* strain used in our study and the strain used in 1982.

The coverage of P22 derived reads showed a distinct pattern of peaks that rise vertically on one side and decline slowly over several 100 kbp increments on the other side. We speculate that the vertical edge of the peak corresponds to the location of the *pac* site at which the packaging of DNA into phage heads is initiated and that the slope of the peak indicates the range of processivity of the headful packaging mechanism (i.e. how many headfuls are packaged into particles before the packaging apparatus dissociates from the chromosome). This speculation is based on several facts: (1) the size of host DNA carried by transducing particles corresponds to the size of the P22 genome (~44 kbp)(31); (2) the P22 genome is replicated by rolling circle replication, which produces long concatemers of P22 DNA. A specific sequence on the phage DNA (*pac* site) initiates the packaging of these concatemers into phage heads using a headful mechanism(31); (3) the packaging of phage DNA continues sequentially along the P22 genome concatemer with a decreasing probability for each next headful to be encapsulated in a phage particle(30); (4) there are five to six sequences on the *S. enterica* genome that are similar to the *pac* site, which leads to packaging of *Salmonella* DNA into P22 particles upon P22 infection, albeit with much lower frequency as compared to P22 DNA(30).

For *E. coli* phage P1, the majority of sequencing reads from purified P1 particles mapped to the P1 genome and only 4.5% of the reads mapped to the *E. coli* genome. The percentage of transducing P1 phages was previously reported to be 6%(32). We also observed that the P1 derived reads mapping to the *E. coli* genome covered the genome unevenly. However, the pattern was less pronounced as compared to P22 and *S. enterica* (Fig. 3c). This low unevenness in sequencing read coverage corresponds to previous data on transduction frequencies of chromosomal markers, which found a maximum transduction frequency across the *E. coli* genome of 10 fold(33).

Sequencing host DNA carried in generalized transducing phages reveals uneven read coverage patterns along the host genome indicative of transduction. These patterns vary in magnitude depending on the transducing phage and they can only be observed if read coverage is analyzed along long stretches of the host genome covering multiples of the length of the DNA carried by the transducing phage e.g. in case of P22 44 kbp. Additionally, the patterns also provide an indication of the frequency with which different regions of the host genome are transduced, as well as the locations of the *pac* sites.

#### Hijacking of helper prophage by a phage-related chromosomal island in *Enterococcus faecalis* and specialized transduction by prophages

Certain chromosomal islands, including pathogenicity islands and integrative plasmids, are mobilized using helper phages(14, 34). This is a form of molecular piracy in which structural proteins of the helper phage are hijacked by the chromosomal island and used as a vehicle for the transfer of the island to other cells. We used *E. faecalis* VE14089, which is a natural resident of the human intestine and causes opportunistic infections, to study the sequencing coverage patterns produced by chromosomal island transfer by way of a helper phage. *E. faecalis* VE14089 is host to a chromosomal island (EfCIV583) that uses structural proteins from a helper phage (pp1) for transfer(15, 34) (Fig. 4a). *E. faecalis* VE14089 possesses five additional prophage-like elements (pp2 to pp6). Some of these prophages contribute to *E. faecalis* pathogenicity and confer an advantage during competition with other *E. faecalis* strains in the intestine(15, 35).

**Figure 4:**
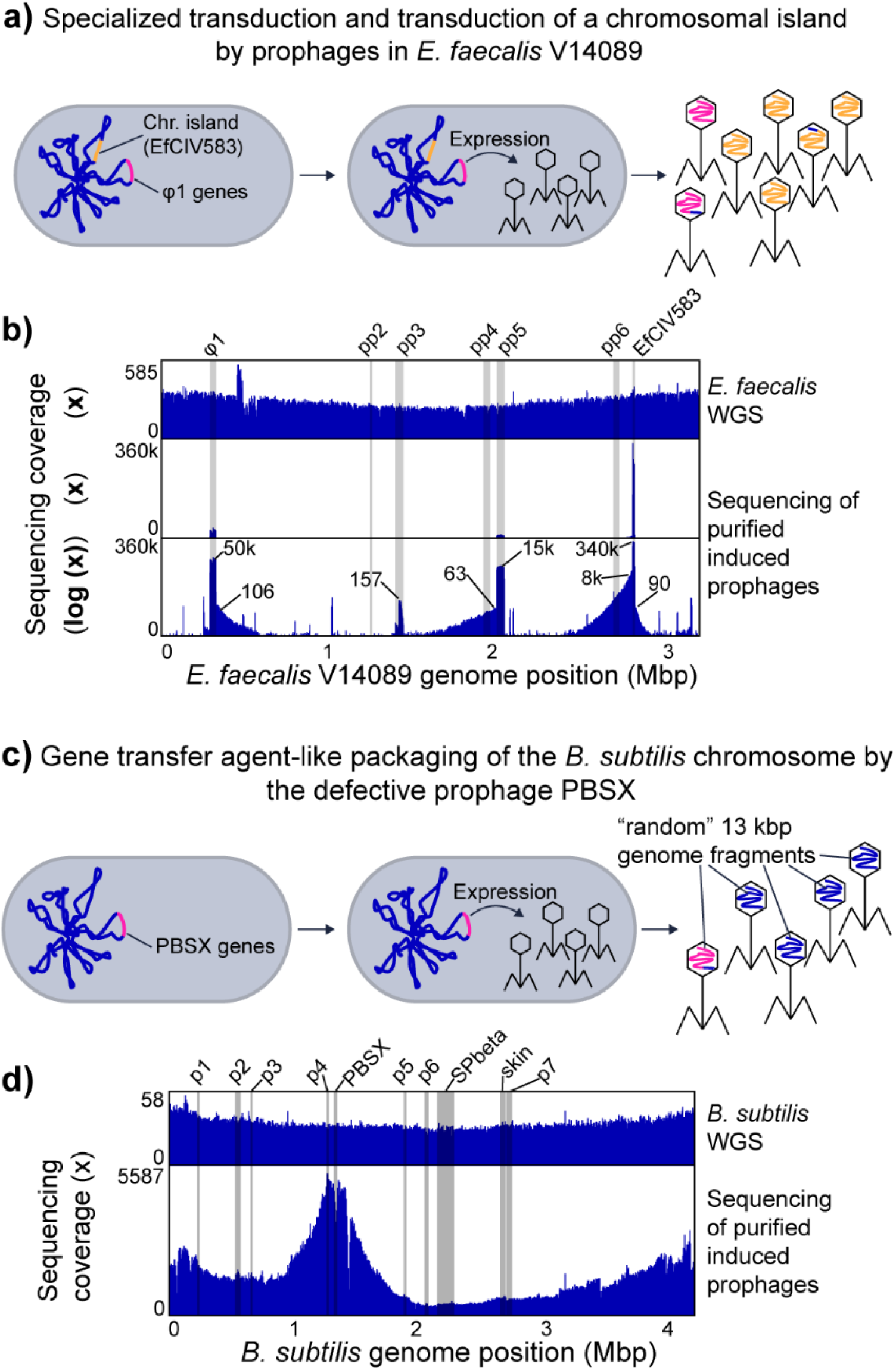
Other types of transduction. a) Specialized transduction (see description for prophage λ) and transduction of a chromosomal island by prophages in *E. faecalis* V14089. The chromosome of *E. faecalis* contains multiple prophages including φ1 and the chromosomal island EfCIV583. Upon induction φ1 and EfCIV583 are excised from the chromosome and replicated. EfCIV583 hijacks the structural proteins of φ1 when they are produced and large number of phage particles that carry the EfCIV583 genome are produced. b) *E. faecalis* V14089 genome coverage patterns associated with prophage induction and EfCIV583 transduction. Whole genome sequencing (WGS) reads and purified VLPs were mapped to the *E. faecalis* genome. The lowest part of the box shows VLP read coverage on a log scale. The small numbers in this plot give x fold coverage for specific genome positions corresponding to prophages or the chromosomal island EfCIV583 and the surrounding areas that are likely transduced. The positions of known prophage-like elements and EfCIV583 in the *E. faecalis genome* are highlighted by grey bars. c) Gene transfer agent-like packaging of the *B. subtilis* chromosome by the defective prophage PBSX. The *B. subtilis* chromosome contains a variety of prophages and prophage-like elements including the defective prophage PBSX(36). Upon expression of the PBSX genes phage-like particles are produced, which contain random 13 kbp pieces of the host chromosome(37). d) *B. subtilis* genome coverage patterns associated with prophage induction. Whole genome sequencing (WGS) reads and purified prophage particle reads were mapped to the *B. subtilis* genome. The positions of known prophages and prophage-like elements in the *B. subtilis* genome (36) are highlighted by grey bars.

Read coverage differs widely between the different prophage like elements with the coverage of EfCIV583 exceeding the coverage of all other elements by almost an order of magnitude (Fig. 4b). This finding is in line with previous results that showed that EfCIV583 DNA is more abundant than all other prophage DNA in purified VLPs from *E. faecalis* V583(35), an isogenic strain of VE14089. Interestingly pp2, pp4 and pp6 did not yield coverage peaks in the VLP fraction, which confirms previous observations that these prophage elements are not excised under the conditions that we used for prophage induction(15).

For pp1, pp5 and EfCIV583 we see patterns (coverage slopes visible in the log-scale coverage plot) that indicate that not only the chromosomal island is transduced but also regions adjacent to these prophages and the chromosomal island. Based on the maximal coverage of the transduced regions versus the coverage of the prophage regions (Fig. 4b) we estimate the maximal frequencies of transduction to be 1:500 for pp1, 1:240 for pp5, and 1:43 for the left side of EfCIV583 and 1:3780 for the right side of EfCIV583. These relatively high transduction frequencies and the fact that transduced regions span several hundred kbp facing unidirectional from the integration site of the prophages and EfCIV583 suggest a lateral transduction mechanism as described by Chen et al.(12).

Our data also revealed that there are several additional regions in the *E. faecalis* VE14089 genome that had an elevated sequencing coverage in the purified phage sample suggesting that these regions encode additional elements that are transported in VLPs. These elements consist of IS-Elements that carry a transposase and surprisingly the three rRNA operons. For the rRNA operons the coverage has a deep valley between the 16S and the 23S rRNA gene suggesting that a specific mechanism for rRNA gene transport is present or that the processed rRNAs were sequenced. We can currently think of three explanations for this intriguing pattern. First, potentially ribosomes are enriched alongside the VLPs in our VLP purification method. However, if this were the case we should have observed similar patterns in VLP fractions of other pure culture organisms, which we did not. Second, intact ribosomes are packaged by VLPs produced in *E. faecalis*. However, this leaves open the question of why the rRNA from these ribosomes was amenable to sequencing by the Illumina method used, which should not enable direct sequencing of RNA. Third, DNA with rRNA genes are packaged with high specificity into one or several types of VLPs from *E. faecalis*.

Our results show that the transduction of a chromosomal island by a prophage can produce a similar coverage pattern as an induced prophage, indicating that chromosomal islands are easy to detect based on read mapping coverage. However, they can only be distinguished from prophage by annotation of the genes and genomic regions.

#### Transport of *Bacillus subtilis* genome by gene transfer agent-like element PBSX

We used induced the gene transfer agent (GTA)-like element PBSX from the *B. subtilis* ATCC 6051 genome to study the sequencing coverage pattern produced by the supposedly randomized incorporation of fragments from the whole genome into GTA type VLPs. PBSX is a defective prophage that randomly packages 13 Kbp DNA fragments of the *B. subtilis* genome in a GTA-like fashion (Fig. 4c)(10, 37, 38). In contrast to other GTAs it does not transfer the packaged DNA between cells but rather acts similar to a bacteriocin against *B. subtilis* cells that do not carry the PBSX gene cluster(10).

DNA sequencing reads derived from purified PBSX particles covered the *B. subtilis* genome unevenly with a maximum 30 fold difference between the lowest and highest covered regions (Fig. 4d). Reads from whole genome sequencing of *B. subtilis* covered the genome evenly slightly increasing toward the origin of replication, as expected(39). The genomic region containing PBSX had a lower read coverage in VLP particle derived reads as compared to neighboring genomic regions. This is consistent with results from a previous study where it was found that a genetic marker integrated in the PBSX region was less frequently packaged into particles as compared to a marker in a neighboring region(40). Interestingly, the genomic region containing the prophage SPbeta, which gets excised upon mitomycin C treatment(41), did not show any higher or lower coverage in the VLP particle derived sequencing reads as compared to neighboring genomic regions (Fig. 4d).

Our results show that packaging of host DNA by the GTA-like PBSX element of *B. subtilis* produces a distinct and non-random sequencing coverage pattern that bears similarities to the read coverage pattern produced by the generalized transducing phage P1 (Fig. 3c).

#### Detection limits of the approach

The patterns for different transduction modes have distinct characteristics that will impact sensitivity of detection and the false positive rate. For prophage induction and specialized transduction pattern detection there are three potential challenges: (1) the length of the genome sequence fragment (contig) used for read mapping needs to be sufficiently long to encompass both the prophage genome, as well as a portion of the host genome; (2) potential assembly artifacts (chimeric contigs consisting of multiple source genomes) can lead to highly uneven read coverage that could look similar to an induced prophage pattern. In the case of our approach this is mitigated by the fact that we map whole metagenome and VLP reads to the same contigs and thus we expect to obtain even read coverage for the whole metagenome read mapping, which is indicative of correct assembly; and (3) if read coverage is too low patterns will not be sufficiently distinct. It can be expected that frequency of specialized transduction is specific to specific host species and prophages. Nevertheless, we tested the lower limit of read coverage levels needed for detection of prophage induction and specialized/lateral transduction by down sampling read numbers for the VLP reads from *E. coli* prophage λ and *E. faecalis* prophages (Figs. 2 and 4b) to achieve coverage levels similar to what we observed for our mouse case study below, which ranged from several tenfold to several thousand fold. For *E. coli* prophage λ we found that at ~6000x maximum read coverage (5% of total reads) the specialized transduction pattern was still weakly visible, but disappeared at lower coverages, while the induction of the prophage itself was still identifiable at read coverages of 20x (0.01% of total reads) and less (Fig. S1a). For the *E. faecalis* prophages specialized/lateral transduction patterns were still visible at ~500x coverage for the pp1 region and at ~150x for the pp5 region. Prophage induction was detectable at coverages well be low 40x (Fig. S1b). These results indicate that specialized and lateral transduction, as well as prophage induction, can sufficiently be detected with read coverages obtained in shotgun metagenomic sequencing of VLPs.

For generalized transduction and GTA mediated DNA transfer pattern detection the two main challenges are; (1) potential generation of similar patterns by contamination of the ultra-purified VLPs with DNA from microbial cells, which can for example be addressed by comparing contig rank abundances between whole metagenome and VLP read coverage (see below in case study); and (2) difficulty to recognize the pattern on short contigs, because sloping can extend across 100s of kbp. To test if generalized transduction or GTA-like patterns can be detected on shorter contigs we used the P22, P1 and PBSX data to simulate how contig length impacts pattern visibility. For this we looked at the coverage patterns of 200 kbp long stretches in the genome (Fig. S2). We found that detectability of generalized and GTA-like patterns in 200 kbp sequence stretches depended on where the 200 kbp stretch was located within the overall read coverage pattern. In some cases distinguishable coverage sloping was observed (e.g. #2 in Fig. S2a and #2 in S2c) in other cases coverage looked even or irregular (e.g. #4 in S2b and #4 in S2c). These results indicate that generalized transduction and GTA mediated DNA transfer can be detected from contig lengths produced using short read shotgun metagenomics of microbiome samples, however, some DNA transfer events are likely missed if the longer contigs do not cover regions that show the characteristic coverage sloping associated with these transfer events.

### Case study: High occurrence of transduction in the intestinal microbiome

We next assessed the power and application of our transductomics approach for detecting transduced DNA in VLPs from complex microbiomes. We sequenced the whole metagenome (~390 mio reads) and VLPs (~360 mio reads) from a fecal sample of one mouse to high coverage. The VLPs were ultra-purified using the multi-step procedure shown in Fig. 1, for which we previously showed that it efficiently removes DNA from microbial cells and the mouse present in fecal samples(24). We were able to assemble 2143 contigs >40 kbp from the whole metagenome reads with the largest contig being 813 kbp (ENA accession for assembly: ERZ1273841). We discarded contigs <40 kbp because detection of transduction patterns requires coverage analysis of a sufficiently large genomic region. We mapped the metagenomic and VLP reads to the contigs >40 kbp to obtain the read coverage patterns. For complete metagenome, 44% of all reads mapped to the contigs >40 kbp and for the VLPs 10% of all reads indicating that a large portion of DNA carried in VLPs is derived from prophages and microbial hosts. Of the 2143 contigs, 1957 showed a “standard” read coverage pattern (Fig. 5a, Suppl. Table S1), i.e. high even coverage of the contigs with metagenomic reads and low even or no coverage with VLP reads, indicating no mobilization of host DNA in VLPs. The remaining 186 contigs (8.6% of all contigs >40 kbp) showed a read coverage pattern that indicates potential mobilization of DNA in VLPs (Fig. 5b-f, Suppl. Table S1).

**Figure 5:**
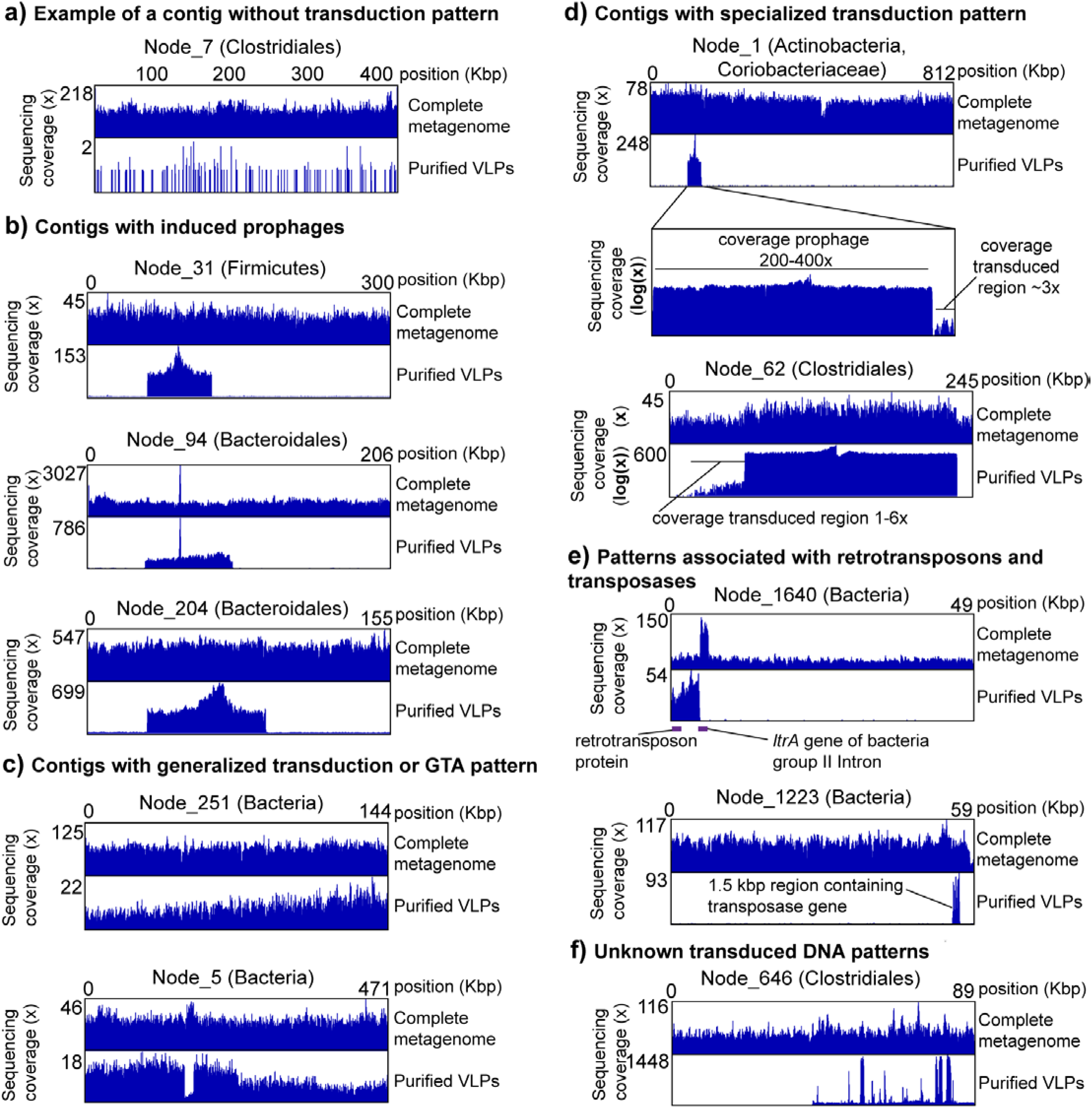
Example of transduction patterns in the mouse intestinal microbiome. The taxonomic classification for each contig specified in parentheses after each contig name is the lowest taxonomic level successfully classified by CAT(42)(Suppl. Table S2). The complete metagenome reads and the purified VLP reads were mapped to the same exact set of contigs assembled from the complete metagenome reads. The read coverage pattern of the complete metagenome reads provides evidence for the correct assembly of the contigs and allows to distinguish potential transduction derived VLP read coverage patterns from VLP coverage patterns due to contamination with microbial DNA. With the exception of panel A) all shown contigs have the same or a lower abundance rank for VLP read coverage as compared to complete metagenome read coverage indicating that their overall read coverage was enriched in the VLP samples. Read coverage due to VLP sample contamination with cellular DNA is expected to result in a higher abundance rank for VLP read coverage, as compared to complete metagenome read coverage.

To verify that the multi-step VLP ultra-purification procedure employed for this study efficiently removes DNA from microbial cells, ruling out potential microbial host DNA contamination, we further assessed read coverage patterns for the 186 contigs in comparison to all contigs. We ranked all 2143 contigs by their normalized coverage for both the whole metagenome and the purified VLP samples (i.e. average x fold read coverage / sum of average x fold read coverage for the sample) with the highest normalized coverage being assigned rank 1 (Table S1). The expectation is that contigs from which DNA is carried in VLPs have the same or lower rank for the VLP sample as compared to the whole metagenome sample, while the rank for contigs for which VLP reads are derived from microbial contamination should have a higher rank as compared to the whole metagenome reads, because randomly contaminating DNA would be depleted in the purified VLP sample. We found that 26 out of the 186 contigs with coverage patterns suggesting DNA mobilization had a normalized coverage based rank that was higher for VLP read coverage than for whole metagenome read coverage indicating that these 26 patterns are potentially due to contamination or alternatively due to very low efficiency of mobilization.

We classified all contigs taxonomically using CAT(42) (Suppl. Tables S2 and S3). The majority of contigs were classified as Bacteroidetes (all contigs: 805, transduction pattern contigs: 83), Firmicutes (all: 586, transduction pattern: 42), Proteobacteria (all: 89, transduction pattern: 3), or not classified at the phylum level (all: 527, transduction pattern: 34). We found that with a few exceptions the relative abundance of contigs assigned to specific phyla was similar between the set of all contigs >40 kbp and the subset of contigs with transduction patterns. The phyla that differed in relative contig abundance were Proteobacteria with less than half the relative abundance in the contigs with transduction patterns, Verrucomicrobia with 3.5x and *Candidatus* Saccharibacteria with 11.5x the contig abundance in the contigs with transduction patterns. Since members of *Cand.* Saccharibacteria have been shown to be extremely small (200 to 300 nm)(43) it is likely that they share similar properties with bacteriophages in terms of size and density and thus might get enriched in the VLP fraction. In fact, all transduction patterns of *Cand.* Saccharibacteria contigs were classified as “unknown” or “unknown, potentially a small bacterium” prior to knowing the taxonomic identity of the contigs.

We classified the type of DNA mobilization/transduction in the 186 contigs with a mobilization pattern based on the visual characteristics of the mobilized region in the VLP read coverage, as well as based on annotated genes within the mobilized region. For example, we classified mobilization patterns as prophage if the characteristic pattern showed high coverage with sharp edges on both sides (compare Fig. 2) and the presence of characteristic phage genes (e.g. capsid proteins) as an additional but not required criterion.

We observed 74 contigs that indicated induced prophages. Of these, 12 (16%) prophages showed indications of specialized transduction i.e. read coverage above the base level of the contig in regions adjacent to the prophage (Fig. 5b and d). Additionally, we classified 8 patterns as potential prophages or chromosomal islands, as they showed the same pattern as other prophages, but we were unable to find recognizable phage genes in the annotations.

We found patterns of potential generalized transduction or GTA carried DNA in 46 contigs, however, some patterns were observed for shorter contigs and could thus potentially be incorrect classifications (Fig. 5c). One of the contigs (NODE_5, classified as Bacteria) with a generalized transduction or GTA pattern additionally showed a sharp coverage drop in a ~15 kbp region only in the VLP reads (Fig. 5c). This region is flanked by a tRNA gene and carries one gene annotated as a potential virulence factor, internalin used by *Listeria monocytogenes* for host cell entry(44). This region might represent a chromosomal island that was excised from the bacterial chromosome prior to or during production of the unknown VLP and that did not get encapsulated in the VLP. Alternatively, similar strains might be present in the sample, but only some carry the chromosomal island and strains carrying the chromosomal island are less prone to producing the VLPs, e.g., by superinfection resistance provided by the chromosomal island against a generalized transducing phage.

We observed 9 patterns that showed strong differences between whole metagenome read coverage and VLP read coverage, but that did not correspond to any of the patterns we analyzed in our proof-of-principle work. However, based on gene annotations we determined that these patterns likely represent retrotransposons or other transposable elements. For example, on contig NODE_1640 (classified as Bacteria by CAT) we observed high coverage with VLP reads on one part of the contig, which carries a gene annotated as a retrotransposon (Fig. 5e). Interestingly, the retrotransposon region is flanked by a ltrA gene which is encoded on bacterial group II intron and encodes maturase, an enzyme with reverse transcriptase and endonuclease activity(45). Surprisingly the region containing the ltrA gene had above average coverage in the whole metagenome reads, but no coverage in the VLP reads. This suggests that the intron actively reverse splices into expressed RNA with subsequent formation of cDNA(45) leading to increased copy number of this genomic region. As another example, on contig NODE_1223 (classified as Bacteria by CAT) a region containing a transposase gene is strongly overrepresented in the VLP reads suggesting that this region is a transposable element that is packaged into a VLP (Fig. 5e).

Finally, we determined that two patterns are likely lytic phages and 47 patterns are classified as “unknown” transduced DNA, as the coverage pattern is uneven indicating transport in VLPs but we could not determine the type of transport. To provide an example, in contig NODE_646 (classified as Clostridiales by CAT) we observed a potential prophage pattern in which we found some of the main phage relevant genes such as major capsid protein, however, within the prophage pattern we observed high coverage spikes for which we currently have no good explanation (Fig. 5f).

## Conclusions and Outlook

The transductomics approach that we developed should be applicable to a broad range of environments ranging from host-associated microbiomes to soils and aqueous environments. For some environments such as open ocean water samples the approach would need only minor modifications. For example, concentration of VLPs by tangential flow filtration prior to density gradient centrifugation using well established protocols(23). Thus this approach will allow addressing key questions about microbial evolution via HGT in a diversity of microbial communities, including what kind of genes and with what frequency are carried by VLPs. Among these questions, one of the most pressing ones is what the role of transducing particles is in the transfer of antibiotic resistance genes, which is a topic of current debate(46–49). Apart from its application in studying transduction in microbial communities this approach can also be used to increase our understanding of the molecular mechanisms of different transduction mechanisms by careful analysis of read coverage data from pure cultures that shows exact transduction frequencies of each genomic location without tedious analysis of multiple genomic markers. Mechanisms that can be analyzed include, for example, the identification of *pac* sites in generalized transducers, the size range of transduced genomic loci in specialized transducers(20), and the analysis of how random DNA packaging by GTAs really is(19).

One of the major surprises for us when analyzing the mouse intestinal transductome data was that around one quarter of the transduction patterns that we identified are unknown. These patterns showed even coverage in the whole metagenome reads and strong uneven coverage in the VLP reads (e.g. Fig. 4e), however, we were unable to associate them clearly with any of transduction modes that we have investigated with pure cultures. We foresee two types of future studies to characterize the nature of the transducing particles that lead to these unknown patterns and to exclude that they are some kind of artifact. First, read coverage patterns of newly discovered modes of transduction have to be analyzed with the “transductomics” approach to correlate the patterns to patterns observed in microbiomes and microbial communities. While we investigated the transduction patterns associated with both major known transduction pathways, as well as more recently discovered transduction pathways, novel modes of transduction are continuously discovered. These novel transduction modes that need to be characterized with our approach include new types of GTAs(10), lateral transduction(12) and DNA transfer in outer membrane vesicles(50, 51). Second, approaches that allow linking specific transduced DNA sequences to the identity of transducing particles in microbial community samples can be developed. We envision, for example, that high resolution filtration and density gradient based separation of individual VLPs will allow linking the transduced DNA (by sequencing) to the identity of the transducing VLPs using proteomics to identify VLP proteins. Using and developing these approaches further will allow us to increase the range of transduction modes that can be detected in microbial communities, as well as potentially reveal currently unknown types of transduction that are not known from pure culture studies yet.

We see several pathways for improving the sensitivity, accuracy and throughput of the transductomics approach in the future. Currently, our ability to detect generalized transduction patterns is limited by the fact that detection of these patterns requires long stretches of the microbial host genome to be assembled. Our P22 and P1 data shows these patterns stretch across genomic regions >500 kbp. Additionally, high sequencing coverage is needed for the detection of these patterns. Assembly of long contigs in metagenomes of high diversity communities is currently hampered by the relatively short read lengths of sequencers that allow for high coverage. We expect, that increasing read numbers of long-read sequencing technologies such as PacBio and Oxford Nanopore in the future will allow us to sequence complex microbiomes to sufficient depth for the assembly of long metagenomic contigs. A combination of long-read sequencing for the whole community metagenomes in combination with a short-read, high-coverage approach for the VLP fraction will in the future provide more sensitive and accurate detection of generalized transduction patterns. In addition to improvements in the realm of long-read sequencing we expect the development of computational tools for the automatic or semi-automatic detection of transduction patterns in read coverage data from paired whole metagenome and VLP metagenome sequencing. There is a large number of possible parameters that could be used to train a machine learning algorithm to detect transduction patterns. These parameters include differences between average read coverage and maximal read coverage for VLP reads (Table S1) and the comparison of contig rank abundance based on coverage, which we used to cross check transduction patterns for signatures of microbial DNA contamination. Such computational tools will enable the high-throughput detection of transduction patterns in many samples, which is currently limited by the need for visual inspection of patterns.

## Online Methods

### *In vitro* bacteriophage propagation and induction of transducing prophages and other elements

#### Lambda

*E. coli* KL740 was inoculated into 300 ml of LB and grown to on OD_600_ of 0.7 at 28ºC with aeration. The culture flask was transferred to a 42ºC water bath for 10 minutes and then incubated at 42º C for 30 min with shaking. The temperature was reduced to 28º C and cell lysis was allowed to proceed for 2 hrs. The remaining cells and debris were removed by centrifugation at 2750 x g for 10 minutes and the phage containing culture fluid was filtered through a 0.45 μm membrane.

#### P22

The data set used to analyze generalized transduction by *Salmonella* phage P22 was taken from a previous study assessing methods for phage particle purification from intestinal contents(24). For a detailed description of P22 propagation and purification please refer to our previous publication.

#### P1

Lyophilized phage P1 was purchased form ATCC and resuspended in 1 ml of Lennox broth (LB) at room temperature (RT). 200 μl of the phage suspension was added to 10 ml of mid logarithmic phase (OD_600_ ~0.5) *E. coli* ATCC 25922 and incubated for 3 hrs at 37ºC with shaking. The bacteria were pelleted at 2750 x g for 10 min and the culture fluid was filtered through a 0.45 μm syringe filter. 100 μl of stationary phase *E. coli* ATCC 25922 was distributed to 15 separate tubes each containing 200 μl of the P1 culture filtrate. The bacteria/phage mixtures were immediately added to molten LB top agar (0.5% agar), poured over LB agar (1.5% agar) plates and incubated overnight (O/N) at 37ºC. 2.5 ml of SM-plus buffer (100 mM NaCl, 50 mM Tris·HCl, 8 mM MgSO_4,_ 5 mM CaCl_2_·6H_2_O, pH 7.4) was added to the surface each plate and the top agar was scraped off and pooled. The phages were eluted from the top agar by rotation for 1 hour at RT. The top agar suspension was centrifuged at 2750 x g for 10 min, the supernatant was collected and the top agar was washed once with ~30 ml of SM-plus and incubated at RT for an additional 1 hr. Following the wash step centrifugation was repeated and the resulting supernatant was collected. The phage containing supernatants were combined.

#### PBSX

To induce the prophage-like element PBSX from the *B. subtilis* ATCC 6051 genome, a 100 ml culture of *B. subtilis* was grown in LB at 37ºC with shaking to an OD_600_ of ~0.5. Mitomycin C was added at a final concertation of 0.5 μg/ml and the culture was incubated at 37ºC for 10 minutes. The culture was centrifuged at 2750 x g for 10 min and the pellet was washed with 50 ml of fresh LB and centrifuged a second time. The cell pellet was resuspended in 100 ml of LB and grown for an additional 3 hrs at 37º C with shaking. The cells and debris were removed by centrifugation and the phage containing culture fluid was filtered through a 0.45 μm membrane.

#### Enterococcal prophages

*E. faecalis* strain VE14089, a derivative of *E. faecalis* V583 that has been cured of its three endogenous plasmids(15), was subcultured to an OD_600_ of 0.025 in 1 L of pre warmed brain heart infusion broth (BHI) and grown statically at 37ºC to an OD_600_ of 0.5. To induce excision of integrated prophages, ciprofloxacin was added to the culture at a final concertation of 2 μg/ml and the bacteria were grown for an additional 4 hrs at 37ºC. The bacterial cells and debris were centrifuged at 2750 x g for 10 min and the culture fluid was filtered through a 0.45μm membrane.

### Purification of phage particles from culture fluid

All phage containing culture fluid was treated with 10 U of DNase and 2.5 U of RNase for 1 hr at RT. 1 M solid NaCl and 10 % wt/vol polyethylene glycol (PEG) 8000 was added and the phages were precipitated O/N on ice at 4ºC. The precipitated phages were resuspended in 2 ml of SM-plus and loaded directly onto CsCl step gradients (1.35, 1.5 and 1.7 g/ml fractions) and centrifuged for 16 hrs at 83,000 x g. The phage bands were extracted from the CsCl gradients using a 23-gauge needle and syringe, brought up to 4 ml with SM-plus buffer and loaded onto a 10,000 Da molecular weight cutoff Amicon centrifugal filter (EMD Millipore) to remove excess CsCl. The phages were washed 3 times with ~4 ml of SM-plus and then stored at 4ºC.

### Isolation of phage and host bacterial DNA from pure cultures

Following CsCl purification of phages and phage-like elements, DNA was isolated by adding 0.5 % SDS, 20 mM EDTA (pH=8) and 50 μg/ml Proteinase K (New England Biolabs) and incubating at 56ºC for 1 hour. Samples were cooled to RT and extracted with an equal volume of phenol:chloroform:isoamyl alcohol. The samples were centrifuged at 12,000 x g for 1 min and the aqueous phase containing the DNA was extracted with an equal volume of chloroform. Following centrifugation at ~16,000 x g for 1 min 0.3M NaOAc (pH=7) was added followed by an equal volume of 100% isopropanol to precipitate the DNA. The DNA was pelleted at 12,000 x g for 30 min and washed once with 500 μl of 70% ethanol. The samples were decanted and the pellets were air dried for 10 min and resuspended in 100 μl of sterile water.

For the isolation of bacterial genomic DNA, we used the Gentra Puregene Yeast/Bacteria Kit (Qiagen) according to the manufacturer’s instructions.

### Isolation and purification of bacteria and VLPs from mouse fecal pellets for metagenomic sequencing

The entire colon contents of one male C57BL6/J mouse were added to 1.2 ml of SM-plus buffer and homogenized manually with the handle of a sterile disposable inoculating loop. After homogenization the sample was brought up to 2 ml with SM-plus. One third of the sample volume was added to a fresh tube containing 100 mM EDTA and set aside on ice. This represented the unprocessed whole metagenome sample. The remaining two thirds of the sample volume were used to isolate VLPs.

VLPs from the homogenized feces were ultra-purified as described previously(24). Briefly, the sample was centrifuged at 2500 x g for 5 min, the supernatant transferred to a clean tube and centrifuged a second time at 5000 x g to pellet any residual bacteria and debris. The supernatant was transferred to a sterile 1 ml syringe and filtered through a 0.45 μm syringe filter. The clarified supernatant was treated with 100 U of DNase and 15 U of RNase for 1 hr at 37ºC. The sample was loaded onto a CsCl step gradient (1.35, 1.5 and 1.7 g/ml fractions) and centrifuged for 16 hrs at 83,000 x g. The VLPs residing at the interface of the 1.35 and 1.5 g/ml fractions were collected (~2 ml) and the CsCl was removed by centrifugal filtration as described above. The purified VLPs were disrupted by the addition of 50 μg/ml proteinase K and 0.5% sodium dodecyl sulfate (SDS) at 56º C for 1 hr. The samples were cooled to room temperature and total DNA was extracted by the addition of and equal volume of phenol:chloroform:isoamyl alcohol. The organic phase was separated by centrifugation at 12,000 x g for 2 minutes and the aqueous phase was extracted with an equal volume of chloroform. The DNA was precipitated by the addition of 0.3 M NaOAc, pH 7, and an equal volume of isopropanol. The DNA pellet was washed once with ice cold 70% ethanol and resuspended in 100 μl of sterile water. The DNA was further cleaned on a MinElute spin column (Qiagen) and eluted into 12 μl of elution buffer (Qiagen).

To purify total metagenomic DNA, unclarified fecal homogenate was treated with 5 mg/ml lysozyme for 30 min at 37º C. The sample was transferred to 2 ml Lysing Matrix B tubes (MP Biomedical) and bead beat in a Bullet Blender BBX24B (Next Advance) at top speed for 1 min followed by placing on ice for 2 min. This was repeated a total of 4 times. The samples were centrifuged at 12,000 x g for 1 min and the DNA from the supernatant was extracted, precipitated and purified as described above.

### DNA Sequencing

The concentration of purified microbial DNA was determined using a Qubit 3.0 fluorometer (Thermo-Fisher). Prior to library preparation total microbial DNA was sheared using a Covaris S2 sonicator with a duty cycle of 10%, intensity setting of 5.0 and a duration of 2 x 60 sec at 4º C. Sequencing libraries were generated using the KAPA HTP library preparation kit KR0426 – v3.13 (KAPA Biosystems) with Illumina TruSeq ligation adapters. Library quality was determined using a 2100 Bioanalyzer system (Agilent). Libraries were size selected and purified in the range of 300-900 bp fragments and subjected to Illumia deep sequencing. For DNA sequencing of pure phage cultures and the *E. coli* KL740 genome we used an Illumina NextSeq 500 desktop sequencer. Illumina Hiseq 2500 v3 Sequencing System in rapid run mode was used to sequence the metagenomes and viromes from the feces of the C57BL6/J mouse. All sequencing was performed in paired end mode acquiring 150 bp reads. Per fragment end the following number of reads were obtained; C57BL/6 mouse feces - 76 M reads for the whole metagenome and 97 M reads for the virome, 45 M reads for phage P1, 21 M reads for lambda phage and 27 M reads for the *E. coli* KL740 genome, 29 M reads for phage PBSX and 28 M reads for the enterococcal prophages. For the C57BL/6 mouse microbiome we sequenced an additional 75 bp single-end read run to increase coverage. We obtained 313 M 75 bp reads for the whole metagenome and 262 M 75 bp reads for the virome. All DNA sequencing was performed by the Eugene McDermott Center for Human Growth and Development Next Generation Sequencing Core Facility at the University of Texas Southwestern.

### Assembly of mouse fecal metagenome

Read decontamination and trimming of the mouse metagenome 75 and 150 bp reads were performed using the BBMap short read aligner (v. 36.19)(25) as previously described(52). Briefly, for decontamination, raw reads were mapped to the internal Illumina control phiX174 (J02482.1), the mouse (mm10) and human (hg38) reference genomes using the bbsplit algorithm with default settings. The resulting unmapped reads were adapter trimmed and low-quality reads and reads of insufficient length were removed using the bbduk algorithm with the following parameters: ktrim = lr, k = 20, mink = 4, minlength = 20, qtrim = f. The reads were assembled using SPAdes version 3.6.1(53) with the following parameters: --only-assembler -k 21,33,55,77,99. Assembled contigs <40 kbp were discarded. The assembly resulted in 2143 contigs >40 kbp.

### Taxonomic classification and annotation of metagenomic contigs

The 2143 contigs >40 kbp from the assembly of the mouse fecal metagenome were taxonomically classified using the Contig Annotation Tool (CAT)(42) (version 2019-07-19). Genes were predicted and annotated using the automated PROKKA pipeline (v1.11) (54).

### Read mapping and read coverage visualization

The whole (meta)genome and purified VLP read sets were mapped onto the corresponding reference genomes of pure culture organisms or the mouse fecal metagenome contigs using BBmap(25) with the following parameters: ambiguous=random qtrim=lr minid=0.97. The generated read mapping files (.bam) were sorted and indexed using SAMtools(v. 1.7)(55). Integrative Genomics Viewer (IGV, v. 2.3.67) tools were used to generate tiled data files (.tdf) from the read mapping (.bam) files for data compression and faster access in IGV using the following parameters: count command, zoom levels: 9, using the mean, window size: 25 or 100(26). Read coverage patterns were displayed and visually assessed in IGV using a linear or if needed log scale.

## Data availability

All sequencing read data generated for this study is available from the European Nucleotide Archive (ENA) through study PRJEB33536 (https://www.ebi.ac.uk/ena/data/view/PRJEB33536). This data includes the reads for the mouse whole metagenomes and VLP fraction, *E. faecalis* VLPs, *B. subtilis* VLPs, *E. coli* phage P1, *E. coli* phage λ, the whole genome sequencing of *E. coli* KL740 (*E. coli* with lambda phage integrated) and the contigs >40 kbp from the mouse whole metagenome assembly (contig file accession number ERZ1273841).

In addition to the de novo generated data we used existing genome assemblies and sequencing read sets for individual bacteria including *Bacillus subtilis* subsp. subtilis str. 168 complete genome from NCBI RefSeq (NC_000964.3), *B. subtilis* 168 genome sequencing read set from the ENA (Study: PRJDB1076, Sample: SAMD00008600), *Enterococcus faecalis* V583 complete genome from NCBI RefSeq (NC_004668.1), *E. faecalis* V583 sequencing read set from the ENA (Study: PRJEB13005, Sample: ERS1085927), *Escherichia coli* K12 complete genome from NCBI RefSeq (NC_000913.3), *E. coli* NCM3722 (*E. coli* K12 with Lambda phage integrated) complete genome sequence from NCBI GenBank (CP011495.1), phage P1 complete genome sequence from NCBI RefSeq (NC_005856.1), *E. coli* K12 genome sequencing read set from the ENA (Study: PRJNA251794, Sample: SAMN02840692), *Salmonella enterica* subsp. enterica serovar typhimurium str. LT2 complete genome sequence from NCBI RefSeq (NC_003197.1), *S. typhimurium* LT2 genome sequencing read set from ENA (Study: PRJNA203445, Sample: SAMN02367645), and a read set of CsCl density gradient purified P22 phage (Study: PRJEB6941, Sample: SAME2690949)(24).

## Author Contributions

M.K. and B.A.D. designed the study. M.K. and B.A.D. performed experiments. M.K. and B.A.D. performed bioinformatic analyses. B.B. developed BBTools and implemented new parameters in BBMap needed for analyses performed in this study. K.S. and L.V.H. provided conceptual input, strains and specialized reagents. M.K. and B.A.D. wrote the paper with input from all authors.

## Acknowledgements

We would like to thank Kelly Ruhn for assistance with animals and Vanessa Schmid and Rachel Bruce of the University of Texas Southwestern Medical Center’s Next Generation Sequencing Core for assistance with Illumina library construction and sequencing support.

This work was supported in part by the USDA National Institute of Food and Agriculture Hatch project 1014212 (M.K.), the Foundation for Food and Agriculture Research Grant ID: 593607 (M.K.), the NC State Chancellor’s Faculty Excellence Program Cluster on Microbiomes and Complex Microbial Communities (M.K.), National Institutes of Health Grants R01AI141479 (B.A.D.), K01DK102436 (B.A.D), and R01DK070855 (L.V.H.), and the Howard Hughes Medical Institute (L.V.H.).

