## Supplementary material for "Microbial DNA on the move: sequencing based detection and analysis of transduced DNA in pure cultures and microbial communities": Figure S1

**Figure S1:** Detection of specialized transduction, lateral transduction and prophage induction read coverage patterns depending on total read coverage. Total VLP read numbers were randomly downsampled to fractions of the total read number using the reformat.sh tool within the BBTools package.

a) Detection limit for specialized transduction by and induction of *E. coli* prophage λ. Images correspond to Panel c) in Figure 2 and were generated with reads down sampled to 5% and 0.01% of the total VLP reads. As in Figure 2c read coverage is shown on a log-scale.

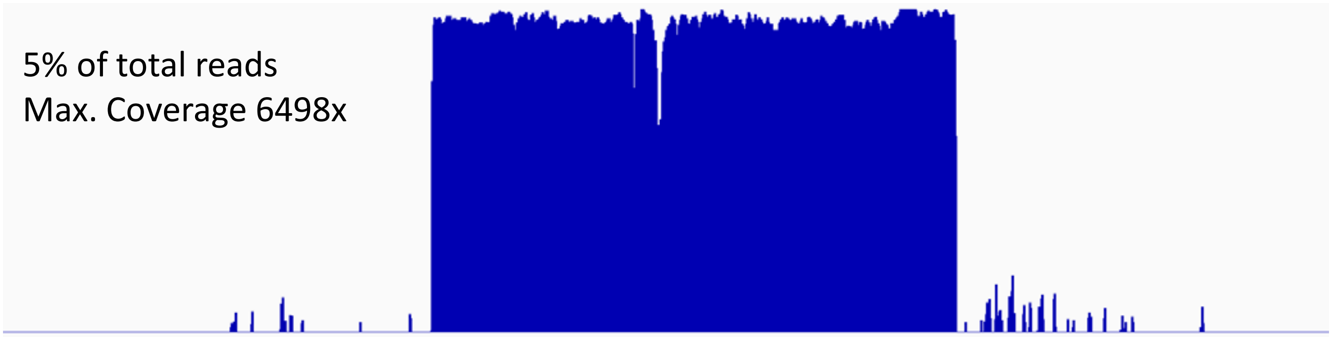


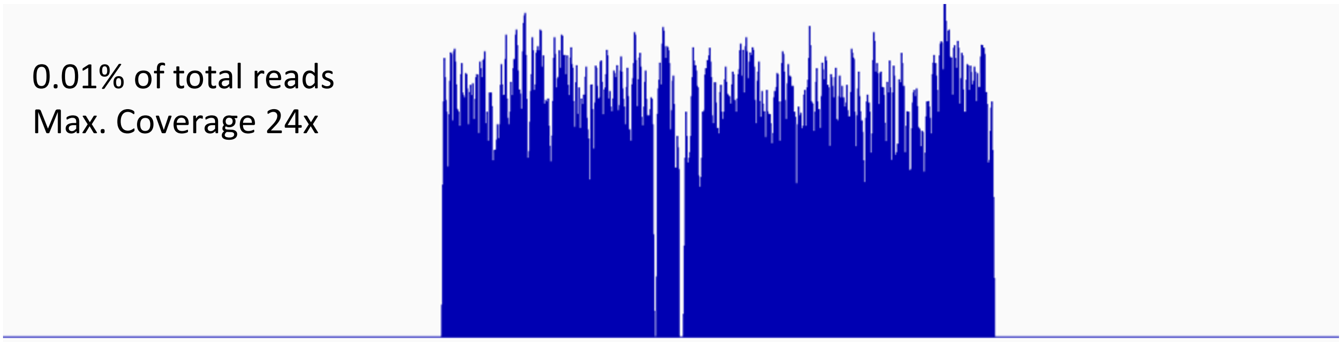

b) Detection limit for lateral transduction by and induction of *E. faecalis* V14089 prophages. Images correspond the bottom of to Panel b) in Figure 4 and were generated with reads down sampled to 1% and 0.01% of the total VLP reads. As in Figure 4b read coverage is shown on a log-scale.

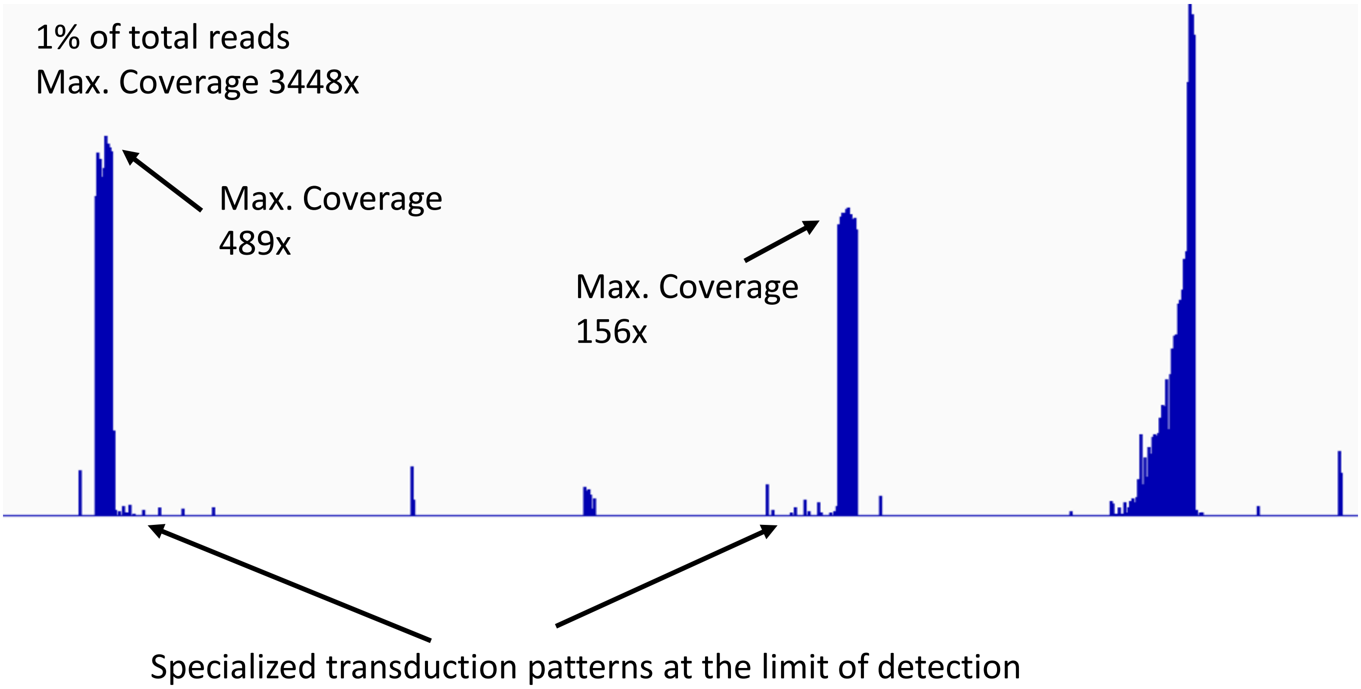


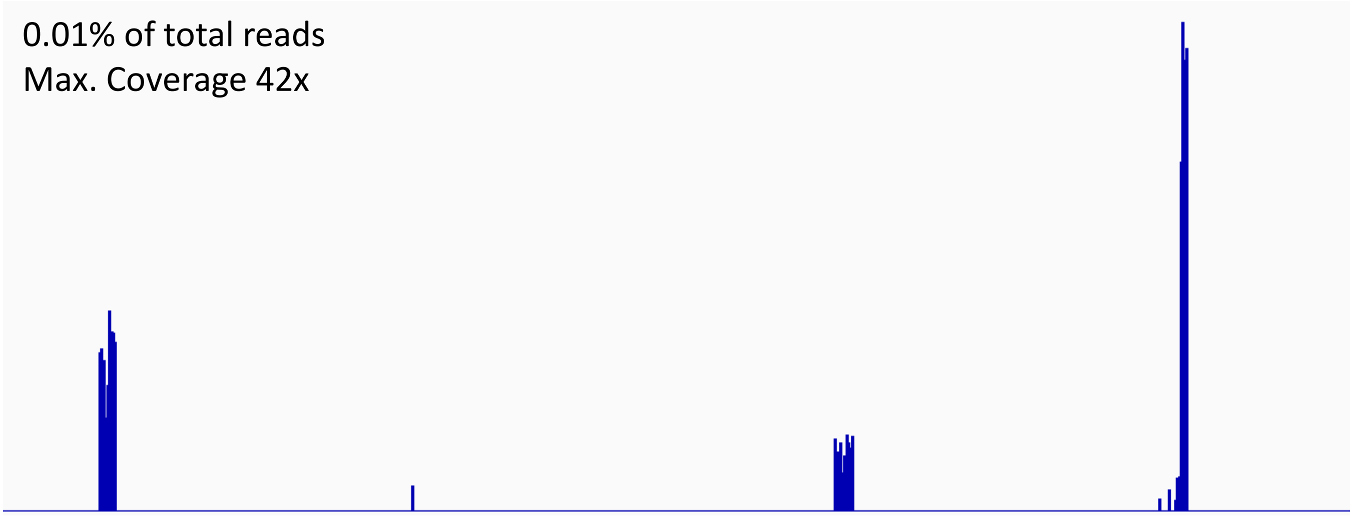
