## Supplementary material for "Microbial DNA on the move: sequencing based detection and analysis of transduced DNA in pure cultures and microbial communities": Figure S2

**Figure S2:** Detection of generalized transduction and GTA-like read coverage patterns depending on contig length. Different contig length were simulated using the P22, P1 and PBSX transduction datasets.

a) Generalized transduction by phage P22

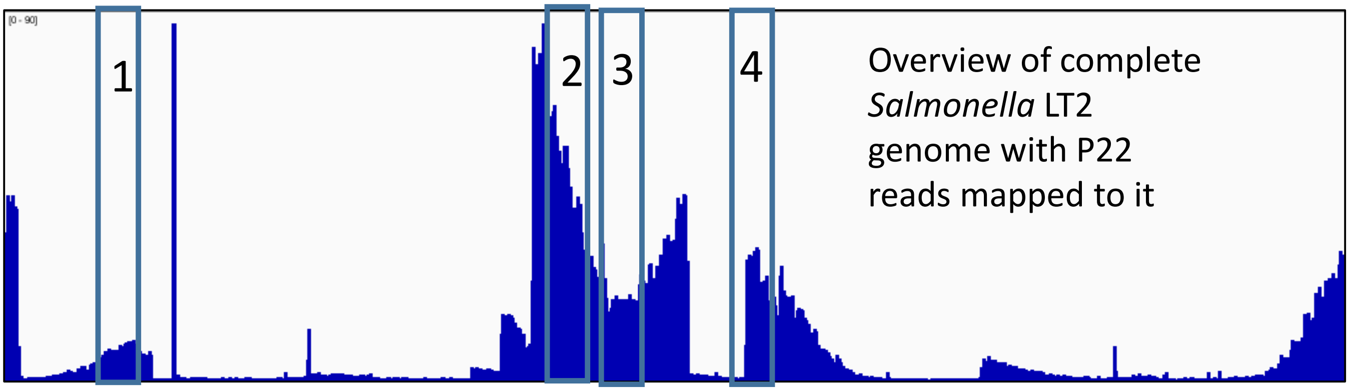


Enlargements of the four 200 kbp sections indicated above showing what the read coverage pattern would look like on a 200 kbp contig for specific regions within the generalized transduction pattern.

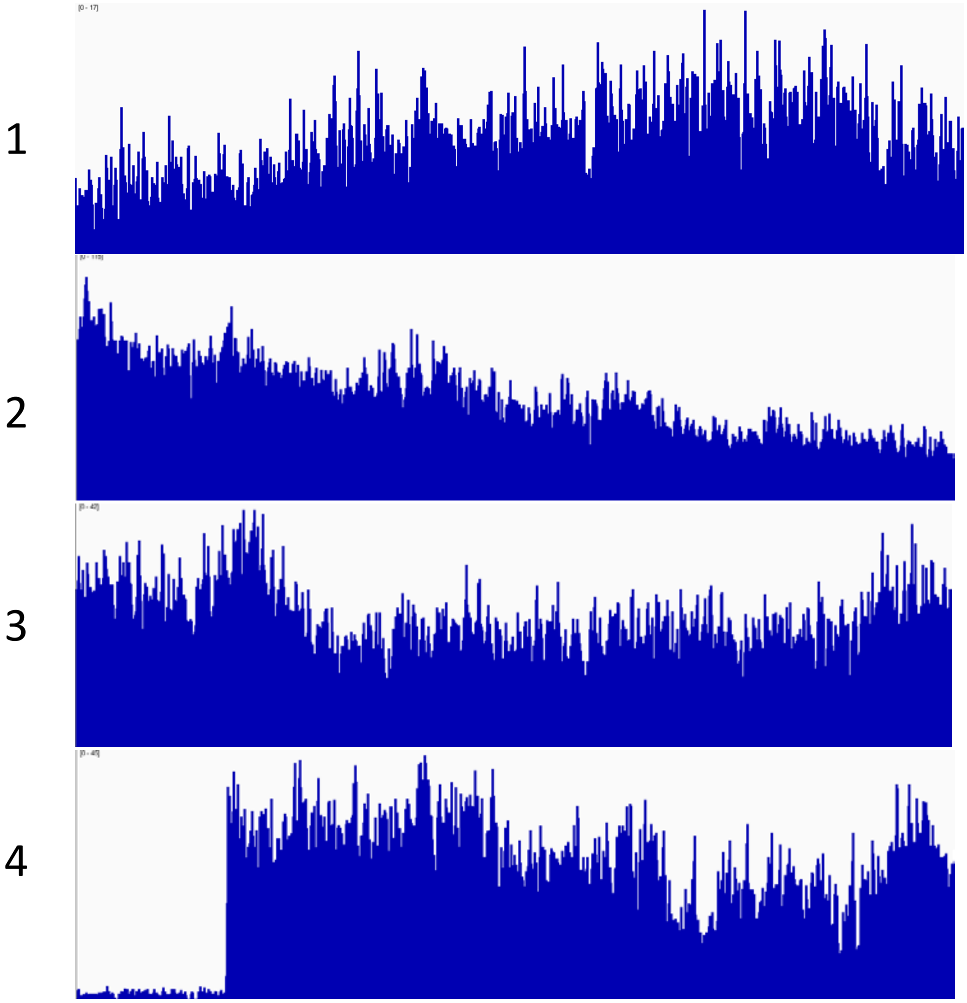

b) Generalized transduction by phage P1

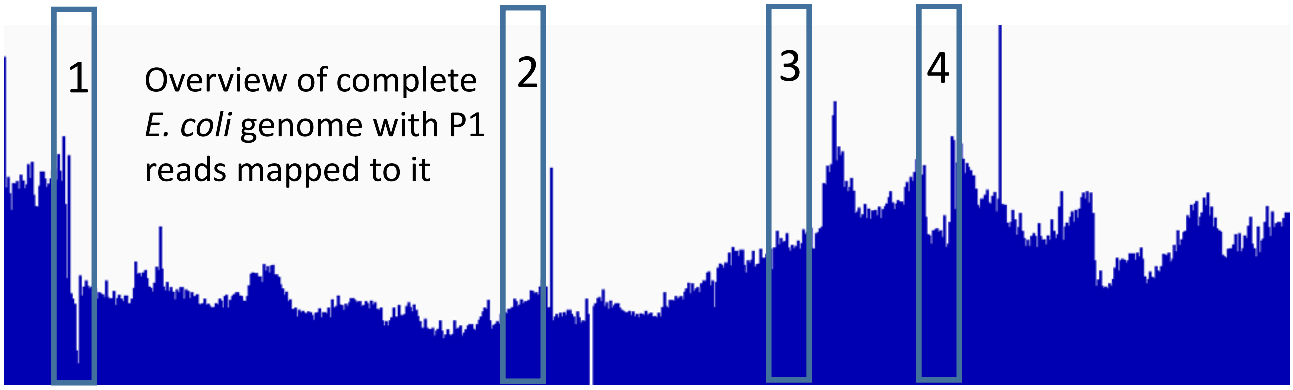


Enlargements of the four 200 kbp sections indicated above showing what the read coverage pattern would look like on a 200 kbp contig for specific regions within the generalized transduction pattern.

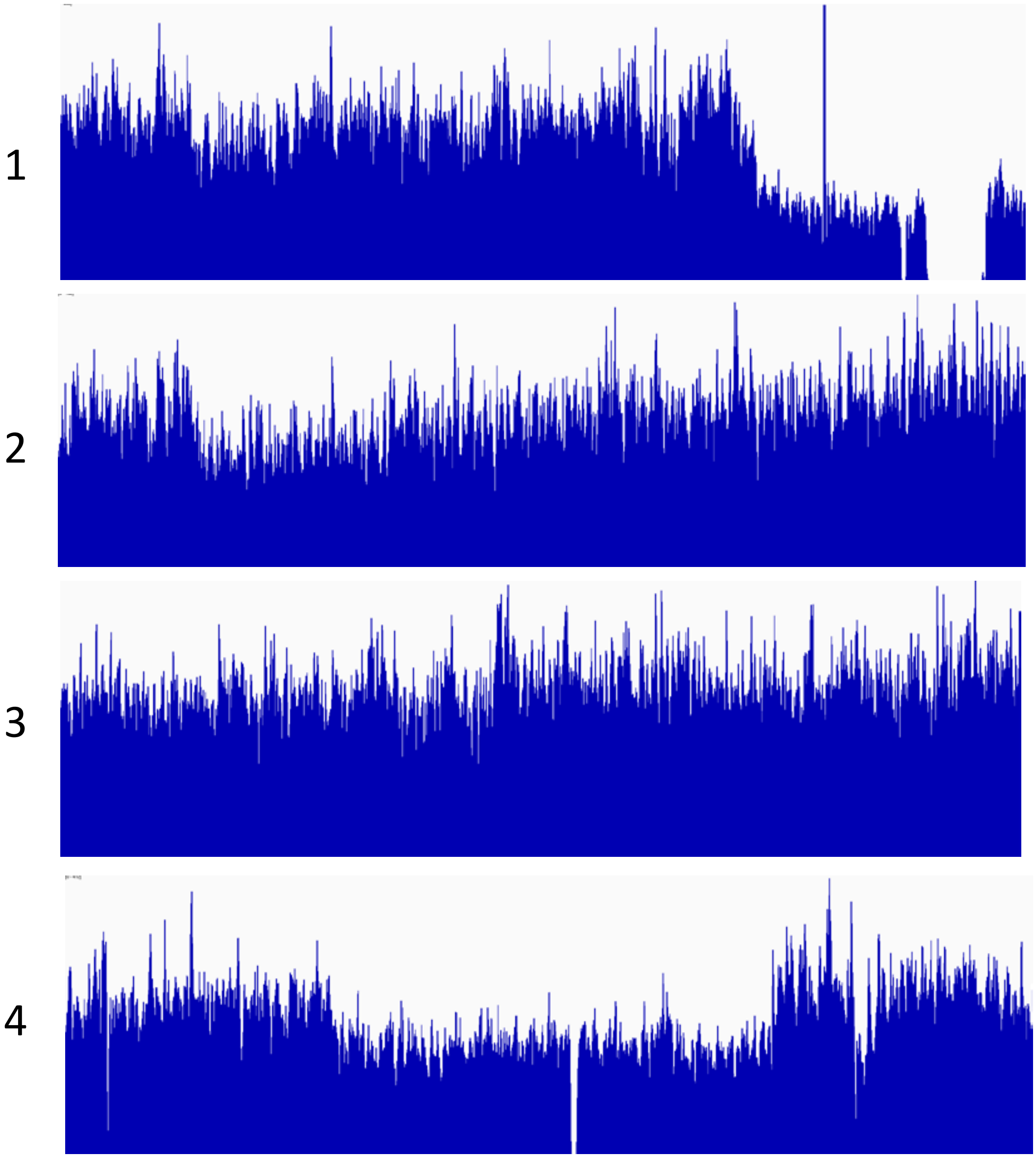

c) GTA-like DNA transport by defective prophage PBSX

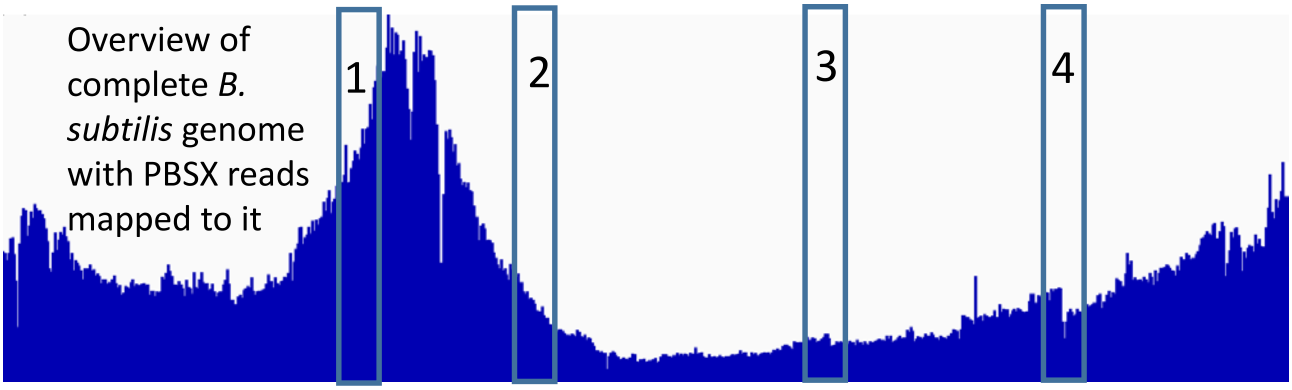


Enlargements of the four 200 kbp sections indicated above showing what the read coverage pattern would look like on a 200 kbp contig for specific regions within the generalized transduction pattern.

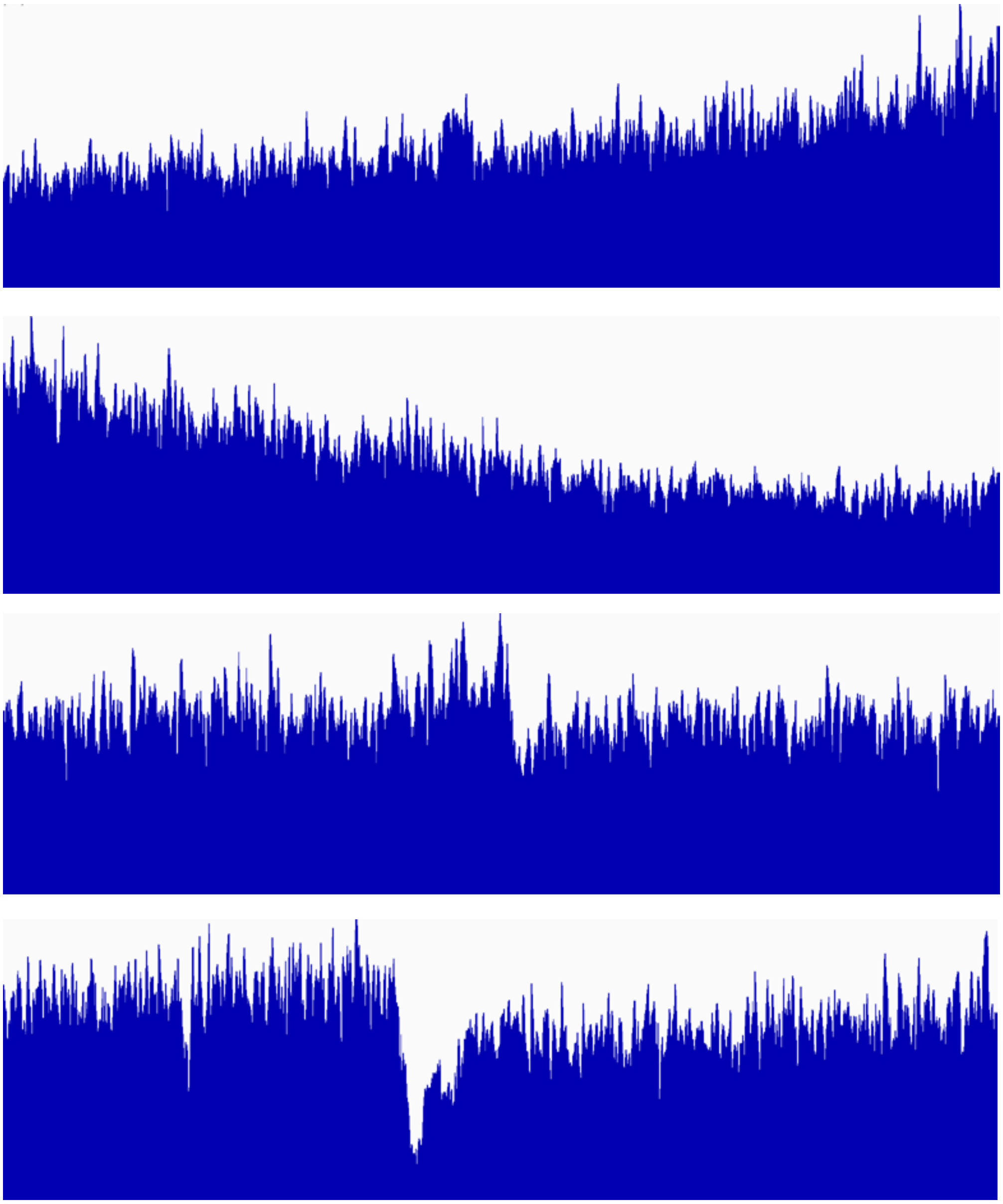

1

2

3

4
